# Why most transporter mutations that cause antibiotic resistance are to efflux pumps rather than to import transporters

**DOI:** 10.1101/2020.01.16.909507

**Authors:** Pedro Mendes, Enrico Girardi, Giulio Superti-Furga, Douglas B. Kell

## Abstract

Genotypic microbial resistance to antibiotics with intracellular targets commonly arises from mutations that increase the activities of transporters (pumps) that cause the efflux of intracellular antibiotics. *A priori* it is not obvious why this is so much more common than are mutations that simply inhibit the activity of uptake transporters for the antibiotics. We analyse quantitatively a mathematical model consisting of one generic equilibrative transporter and one generic concentrative uptake transporter (representing any number of each), together with one generic efflux transporter. The initial conditions are designed to give an internal concentration of the antibiotic that is three times the minimum inhibitory concentration (MIC). The effect of varying the activity of each transporter type 100-fold is dramatically asymmetric, in that lowering the activities of individual uptake transporters has comparatively little effect on internal concentrations of the antibiotic. By contrast, increasing the activity of the efflux transporter lowers the internal antibiotic concentration to levels far below the MIC. Essentially, these phenomena occur because inhibiting individual influx transporters allows others to ‘take up the slack’, whereas increasing the activity of the generic efflux transporter cannot easily be compensated. The findings imply strongly that inhibiting efflux transporters is a much better approach for fighting antimicrobial resistance than is stimulating import transporters. This has obvious implications for the development of strategies to combat the development of microbial resistance to antibiotics and possibly also cancer therapeutics in human.

## Introduction

In order to understand genotypic antimicrobial resistance and how to combat it, a starting point should be an understanding of the main kinds of mutation that can cause it. For present purposes, we assume that the molecular targets of the antibiotic are intracellular (and indeed when the microbes themselves are inside host cells, their access presents its own problems ^1^). Broadly, these mutations are of then of three kinds ^2–4^: (i) mutations in or overproduction of one or more targets of the antibiotic (e.g. DNA gyrase and topoisomerase IV for ciprofloxacin ^5^), (ii) mutations that lead to inactivation of the antibiotic (e.g. of chloramphenicol ^6^ and aminoglycosides ^7^), or (iii) mutations that affect the ability of the antibiotic to be transported to a compartment containing its sites of action in the target microbe.

To enter the target microbe, antibiotics (as do other drugs, e.g. ^8–14^) require transporters. (In Gram-negatives, outer-membrane proteins may also play a role ^15–17^.) The precise identities of these uptake transporters are in general not well understood, because mutations tend to lead only to partial resistance. However, they have been identified for antibiotics such as aminoglycosides ^18^, chloramphenicol ^19^, cycloserine ^20^ and fosfomycin ^21, 22^. In addition, bacteria have also evolved a variety of efflux pumps that serve to remove such antibiotics (see later, and also many other substances ^23, 24^) from the cells. Thus, mutations that affect transporter activity can in principle involve uptake transporters, efflux transporters, or upstream regulators of their activity. Our focus is on this collective class, viz. transporters. In particular, consistent with the difficulty of identifying transporters for their uptake, we note that the very great bulk of transporter-mediated resistance is mediated via (multi-drug) efflux rather than influx transporters (e.g. ^25–45^). The focus of this article is to enquire as to the reasons why this might be so.

To this end, we create a very simple and generic model (Fig 1), consisting of two types of influx and one type of efflux transporter. For the influx transporters, one is a generic equilibrative transporter and one is concentrative for uptake, i.e. it has the capability of raising the concentration of the drug of interest to a higher level inside than outside. Such transporters necessarily require a source of free energy; in prokaryotes this is mainly ATP ^46, 47^. The effluxer is also taken to be ATP-driven. We assume that a drug (antibiotic) has been added at 3x the minimum inhibitory concentration (MIC), which for our purposes is taken to be 1 concentration unit in the case of the wild type, but that the drug does not itself alter the expression levels of the transporters (cf. ^48^).

**Fig 1.**
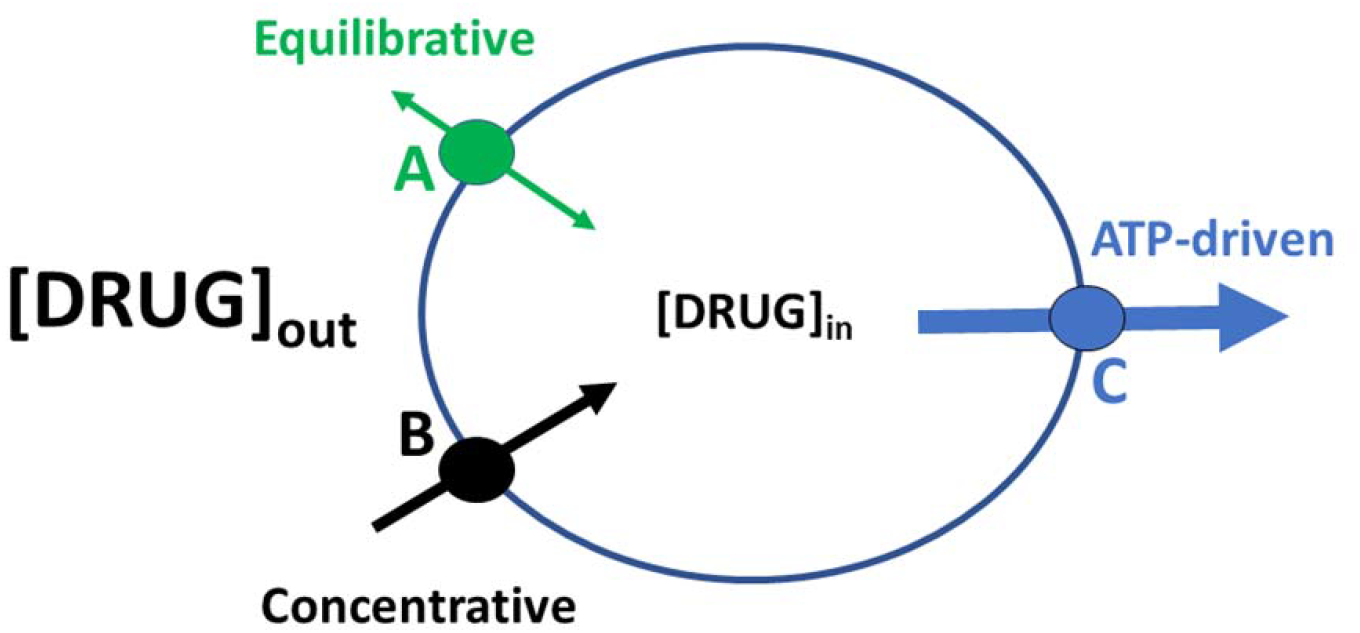
The generic model in which we have a suite of (A) equilibrative and (B) concentrative influx transporters, together with a generic ATP-driven efflux transporter.

Intuitively, lowering the internal concentration of the drug by blocking the concentrative one only works if the equilibrative ones are collectively slower than an individual concentrator, and this is unlikely if there are several. Similarly, trying to lower the internal concentration by blocking one of the equilibrative ones would just let the concentrative one(s) ‘pick up the slack’. This already suggests the general reason why a partial inhibition of uptake activity might have comparatively little effect. Of course if we start with the drug at a level above its MIC it is clear that increasing the effluxer activity can serve to bring to a level below the MIC (and that lowering any starting efflux activity would increase antibiotic sensitivity). We now wish to assess these intuitions by putting some concrete numbers on these fluxes. In systems biology ^49–53^, this is commonly done by casting the enzymatic rate equations into the form of ordinary differential equations, and this is what we do here.

## Materials and methods

As previously ^54^, all simulations were performed using COPASI, here version 4.27, with the LSODA integrator ^55–57^ (http://copasi.org/), which reads and writes SBML-compliant models ^58–60^. It contains a full suite of enzyme rate equations, and admits automated parameter sweeps. Model files including the precise parameters are included as supplementary data.

The simulations were carried out with a differential equation-based model with three compartments (Fig 1), viz. the intracellular space, the inner membrane, and the extracellular space (including the periplasmic volume). Three different transporters are considered: transporter *A* is an equilibrator that allows transport in both directions (*K*_*eq*_ = 1), *B* is a concentrative influx transporter; even though allowing transport in both directions, it favors transport into the cell (modelled by setting *K*_*eq*_ = 10 or *K*_*eq*_ = 100). *C* is an efflux pump that only transports the drug from the cytoplasm to the outside.

The model was set up to mimic typical assays, and parameters were set to values that are comparable to what is found in the literature as follows. Total volume of the assay is 150 μl (from ^61^). Each assay is estimated to have 10^6^ cells, with an average volume of 4×10^−15^ l per cell ^62^ (grown in rich media). Estimates of the proportion of volume taken by the periplasm are around 30% ^63^. Thus, the total cell volume in the assay is estimated at 4×10^−9^ l and the cytoplasmic volume at 2.8×10^−9^ l. For the inner membrane surface area we adopt the average value in the range considered by Wong and Amir ^64^ 34.5 μm^2^ (3.45×10^−7^ cm^2^), which corresponds to a total surface area of 0.345 cm^2^ (*i.e.* for all 10^6^ cells); note that Thanassi *et al.* provide an estimate 3-fold lower (0.103 cm^2^) 65.

Kinetic parameters for the efflux pump (*C*) come from Nagano and Nikaido for AcrB (part of acrAB/tolC) with nitrocefin 66; they cite a *K*_*m*_ of 5 μM, *K*_*cat*_ of 10 s^−1^ and a *V*_*max*_ of 2.35×10^−11^ mol/s/10^9^ cells, which implies a total of 2.35×10^−12^ mol of transporter. Considering that our simulation contains 10^6^ cells, the adjusted amount of transporter is then 2.35×10^−15^ mol (considering the surface area estimated above, this corresponds to a surface density of 6.8×10^−15^ mol/cm^2^) with a *V*_*max*_ of 2.35×10^−14^ mol/s, assuming the same *k*_*cat*_ as for nitrocefin. For *K*_*m*_ we chose a higher value (500 μM).

## Results

Fig 2 shows our ‘baseline simulation, in which a steady-state intracellular level of the drug similar to that outside is obtained by balancing the three main fluxes.

**Fig 2.**
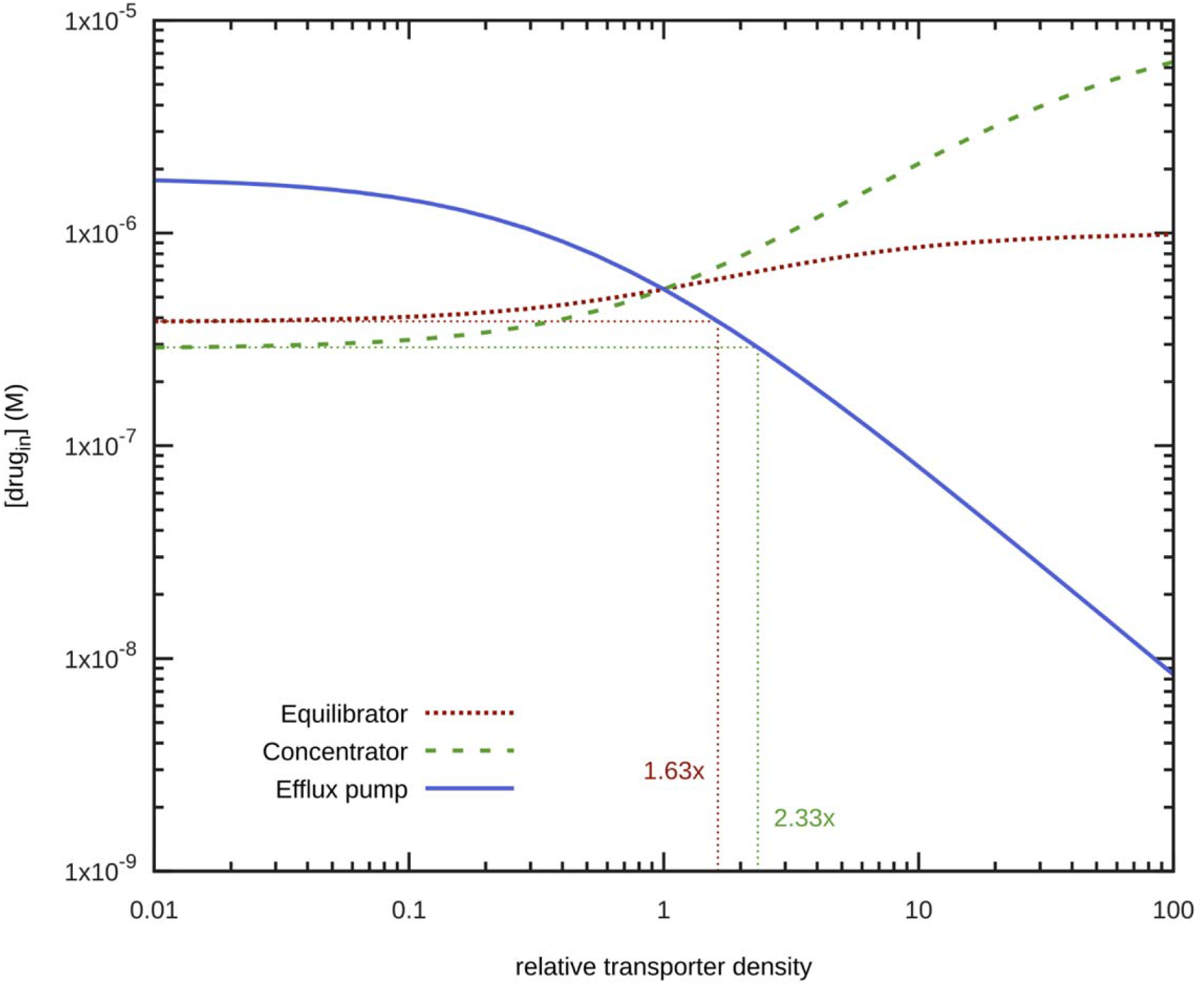
Effect of varying the relative rates of the three generic transporters individually on the normalized accumulation of an antibiotic. Parameters as in Methods and the supplementary files, with K_eq_ for transporter B set at 10.

It is clear that there is a very strong asymmetry; decreasing the individual activities of the equilibrative or concentrative transporters even 100-fold has only a 1.63- or 2.33-fold effect on the steady-state intracellular concentration of the drug, while increasing the effluxer activity by the same amount lowers the intracellular concentration fifty-fold.

Changing the (maximal) degree to which the concentrator concentrates (viz 100-fold rather than 10-fold) also has no material effect on the results when individual transporter activities are lowered, and only a marginal effect when the activity of the concentrator is raised (Fig 3, top right).

**Fig 3.**
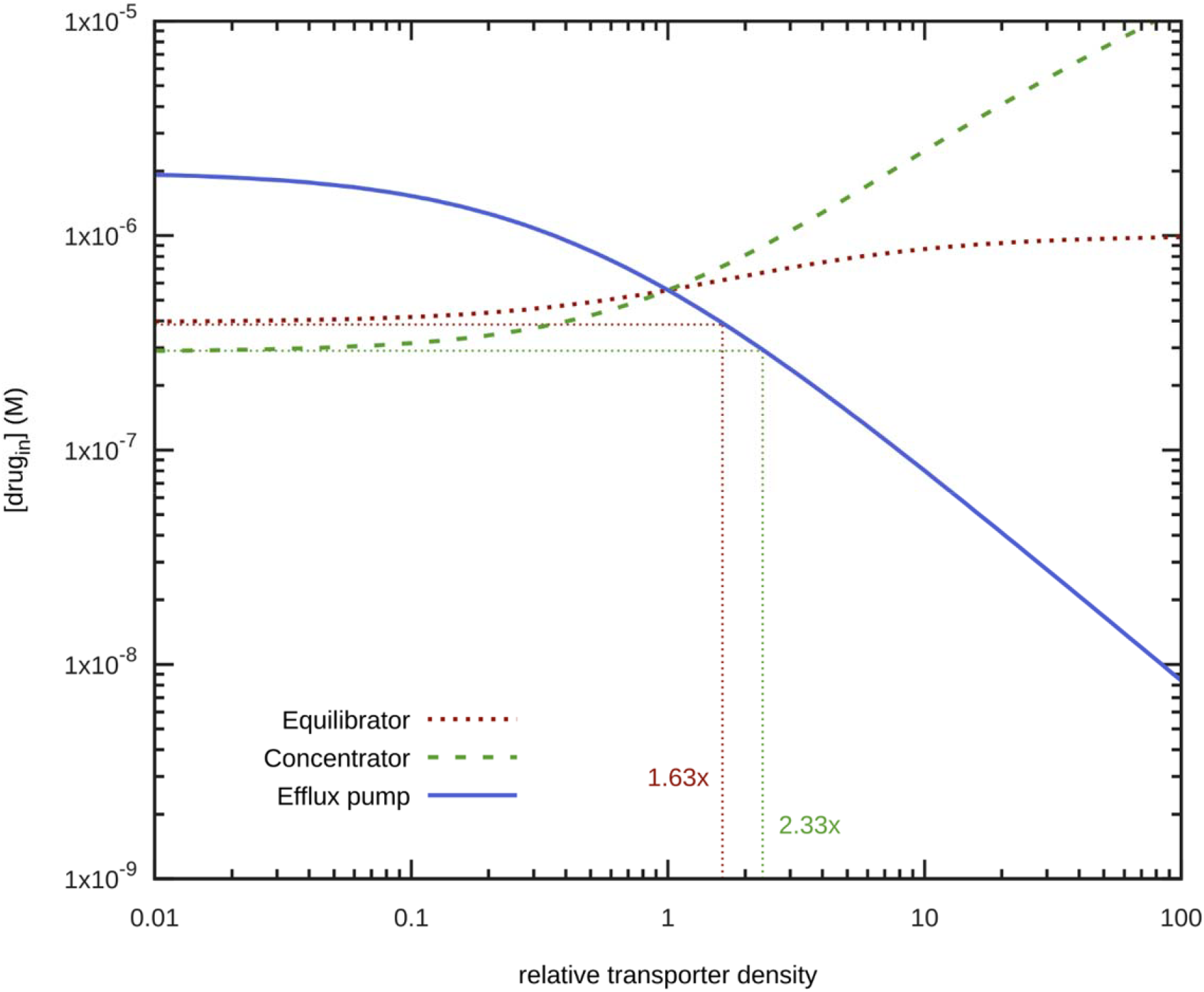
Effect of varying the relative rates of the three generic transporters on the normalized accumulation of accumulation of an antibiotic. Parameters as in Methods and the supplementary files, with K_eq_ for transporter B set at 100.

## Discussion

Microbial resistance to antibiotics (AMR) remains a huge problem (e.g. ^67–72^). To this end, a major cause is the ability of efflux pumps to create resistance to antibiotics by pumping them out from the cytoplasm of cells (e.g. ^25–45^). This is true for cytotoxic substances more generally, including anti-cancer drugs ^42, 48^. Many efflux transporters are sufficiently active that even when the drug has relatively tight intracellular binding sites they can effectively remove almost all of it, as is the case with AcrAB/TolC and ethidium bromide ^73, 74^. A recent experimental survey of several hundred gene knockouts in *E. coli*, using fluorescent probes as antibiotic surrogates showed that dozens of such efflux transporters could be active and thereby contribute to lowering the steady-state uptake ^47^. There is also considerable redundancy and plasticity ^75^. Thus, as expected from metabolic control analysis, while there is little effect of single-gene knockouts on fluxes ^76^, there can be potentially very large effects on the concentrations of intermediary metabolites ^77, 78^ or, as in our model, the intracellular concentration of an antibiotic of interest,

If there is only a single influx transporter (or one that is overwhelmingly dominant) for a cytotoxic drug of interest, as occasionally happens ^13^, inhibiting it can lower the toxicity of the drug enormously; in the case of YM155 (sepantronium bromide) this could be by several hundredfold ^13^. However, it is possible that mutation of a non-redundant influx transporter might also induce significant metabolic costs, although there are also constraints ^79^. Moreover, most cytotoxic drugs can be taken up by multiple transporters ^80, 81^, and affecting all of them simultaneously is probably not realistic.

The consequences of our simple model are thus clear: in order to inhibit the development of antimicrobial resistance, we need to be able to inhibit the efflux pumps that such bacteria possess and use in abundance. To this end, it is indeed widely considered that inhibitors of efflux pumps might well have a role to play in reducing AMR ^42, 82–85^. The present simulations put this thinking on a firm and quantitative footing.

## Supporting information

Model files and data

## Acknowledgements

DBK thanks the BBSRC (grants BB/P009042/1 and BB/R000093/1), the Novo Nordisk Fonden via the Centre for Biosustainability (grant NNF10CC1016517), and the University of Liverpool for financial support. PM thanks the NIH (grants GM115043 and GM127909) for financial support. EG and GSF acknowledge supports from the Austrian Academy of Sciences and the European Research Council (ERC AdG 695214 GameofGates).

## Supplementary information

A zip file containing the COPASI model and results files.

## Author contribution statement

EG and GSF originally posed the problem to DBK. DBK defined a suitable system and suggested the idea of modelling it. PM ran all the simulations. All authors contributed to the writing of the ms.

## References

1. Prideaux, B. et al. The association between sterilizing activity and drug distribution into tuberculosis lesions. Nat Med 21, 1223–1227 (2015).

2. McKeegan, K.S., Borges-Walmsley, M.I. & Walmsley, A.R. Microbial and viral drug resistance mechanisms. Trends Microbiol 10, S8–14 (2002).

3. Strateva, T. & Yordanov, D. *Pseudomonas aeruginosa* - a phenomenon of bacterial resistance. J Med Microbiol 58, 1133–1148 (2009).

4. Munita, J.M. & Arias, C.A. Mechanisms of Antibiotic Resistance. Microbiol Spectrum 4, VMBF-0016–2015 (2016).

5. Rehman, A., Patrick, W.M. & Lamont, I.L. Mechanisms of ciprofloxacin resistance in *Pseudomonas aeruginosa*: new approaches to an old problem. J Med Microbiol 68, 1–10 (2019).

6. Shaw, W.V. et al. Primary structure of a chloramphenicol acetyltransferase specified by R plasmids. Nature 282, 870–872 (1979).

7. Ramirez, M.S. & Tolmasky, M.E. Aminoglycoside modifying enzymes. Drug Resist Updat 13, 151–171 (2010).

8. Dobson, P.D. & Kell, D.B. Carrier-mediated cellular uptake of pharmaceutical drugs: an exception or the rule? Nat Rev Drug Disc 7, 205–220 (2008).

9. Fromm, M.F. & Kim, R.B. (eds.) Drug Transporters, Vol. 201. (Springer, Berlin; 2011).

10. You, G. & Morris, M.E. (eds.) Drug Transporters: Molecular Characterization and Role in Drug Disposition, Edn. 2nd. (Wiley, New York; 2014).

11. Kell, D.B., Dobson, P.D., Bilsland, E. & Oliver, S.G. The promiscuous binding of pharmaceutical drugs and their transporter-mediated uptake into cells: what we (need to) know and how we can do so. Drug Disc Today 18, 218–239 (2013).

12. Kell, D.B. & Oliver, S.G. How drugs get into cells: tested and testable predictions to help discriminate between transporter-mediated uptake and lipoidal bilayer diffusion. Front Pharmacol 5, 231 (2014).

13. Winter, G.E. et al. The solute carrier SLC35F2 enables YM155-mediated DNA damage toxicity. Nat Chem Biol 10, 768–773 (2014).

14. Giacomini, K.M., Galetin, A. & Huang, S.M. The International Transporter Consortium: Summarizing Advances in the Role of Transporters in Drug Development. Clin Pharmacol Ther 104, 766–771 (2018).

15. Hancock, R.E.W. The bacterial outer membrane as a drug barrier. Trends Microbiol 5, 37–42 (1997).

16. Bajaj, H. et al. Antibiotic uptake through membrane channels: role of *Providencia stuartii* OmpPst1 porin in carbapenem resistance. Biochemistry 51, 10244–10249 (2012).

17. Mahendran, K.R., Kreir, M., Weingart, H., Fertig, N. & Winterhalter, M. Permeation of antibiotics through Escherichia coli OmpF and OmpC porins: screening for influx on a single-molecule level. J Biomol Screen 15, 302–307 (2010).

18. Taber, H.W., Mueller, J.P., Miller, P.F. & Arrow, A.S. Bacterial uptake of aminoglycoside antibiotics. Microbiol Rev 51, 439–457 (1987).

19. Prabhala, B.K. et al. The prototypical proton-coupled oligopeptide transporter YdgR from *Escherichia coli* facilitates chloramphenicol uptake into bacterial cells. J Biol Chem 293, 1007–1017 (2018).

20. Chen, J.M., Uplekar, S., Gordon, S.V. & Cole, S.T. A point mutation in cycA partially contributes to the D-cycloserine resistance trait of *Mycobacterium bovis* BCG vaccine strains. PLoS One 7, e43467 (2012).

21. Takahata, S. et al. Molecular mechanisms of fosfomycin resistance in clinical isolates of *Escherichia coli*. Int J Antimicrob Agents 35, 333–337 (2010).

22. Ballestero-Téllez, M. et al. Molecular insights into fosfomycin resistance in *Escherichia coli*. J Antimicrob Chemother 72, 1303–1309 (2017).

23. Kell, D.B., Swainston, N., Pir, P. & Oliver, S.G. Membrane transporter engineering in industrial biotechnology and whole-cell biocatalysis. Trends Biotechnol 33, 237–246 (2015).

24. Kell, D.B. in Fermentation microbiology and biotechnology, 4th Ed. (eds. E.M.T. El-Mansi, J. Nielsen, D. Mousdale, T. Allman & R. Carlson) 117–138 (CRC Press, Boca Raton; 2019).

25. Phillips-Jones, M.K. & Harding, S.E. Antimicrobial resistance (AMR) nanomachines-mechanisms for fluoroquinolone and glycopeptide recognition, efflux and/or deactivation. Biophys Rev 10, 347–362 (2018).

26. Chopra, I. & Roberts, M. Tetracycline antibiotics: Mode of action, applications, molecular biology, and epidemiology of bacterial resistance. Microbiol Mol Biol Rev 65, 232−+ (2001).

27. Alekshun, M.N. & Levy, S.B. Molecular mechanisms of antibacterial multidrug resistance. Cell 128, 1037–1050 (2007).

28. Bhardwaj, A.K. & Mohanty, P. Bacterial efflux pumps involved in multidrug resistance and their inhibitors: rejuvinating the antimicrobial chemotherapy. Recent Pat Antiinfect Drug Discov 7, 73–89 (2012).

29. Nikaido, H. & Pagès, J.M. Broad-specificity efflux pumps and their role in multidrug resistance of Gram-negative bacteria. FEMS Microbiol Rev 36, 340–363 (2012).

30. Blair, J.M., Richmond, G.E. & Piddock, L.J.V. Multidrug efflux pumps in Gram-negative bacteria and their role in antibiotic resistance. Future Microbiol 9, 1165–1177 (2014).

31. Delmar, J.A., Su, C.C. & Yu, E.W. Bacterial multidrug efflux transporters. Annu Rev Biophys 43, 93–117 (2014).

32. Sun, J., Deng, Z. & Yan, A. Bacterial multidrug efflux pumps: mechanisms, physiology and pharmacological exploitations. Biochem Biophys Res Commun 453, 254–267 (2014).

33. Anes, J., McCusker, M.P., Fanning, S. & Martins, M. The ins and outs of RND efflux pumps in *Escherichia coli*. Front Microbiol 6, 587 (2015).

34. Li, X.Z., Plésiat, P. & Nikaido, H. The challenge of efflux-mediated antibiotic resistance in Gram-negative bacteria. Clin Microbiol Rev 28, 337–418 (2015).

35. Hernando-Amado, S. et al. Multidrug efflux pumps as main players in intrinsic and acquired resistance to antimicrobials. Drug Resist Updat 28, 13–27 (2016).

36. Alibert, S. et al. Multidrug efflux pumps and their role in antibiotic and antiseptic resistance: a pharmacodynamic perspective. Expert Opin Drug Met Toxicol 13, 301–309 (2017).

37. Hassan, K.A. et al. The putative drug efflux systems of the *Bacillus cereus* group. PLoS One 12, e0176188 (2017).

38. Yılmaz, Ç. & Özcengiz, G. Antibiotics: Pharmacokinetics, toxicity, resistance and multidrug efflux pumps. Biochem Pharmacol 133, 43–62 (2017).

39. Ahmad, I. et al. Bacterial Multidrug Efflux Proteins: A Major Mechanism of Antimicrobial Resistance. Curr Drug Targets 19 (2018).

40. Du, D. et al. Multidrug efflux pumps: structure, function and regulation. Nat Rev Microbiol 16, 523–539 (2018).

41. Zgurskaya, H.I., Rybenkov, V.V., Krishnamoorthy, G. & Leus, I.V. Trans-envelope multidrug efflux pumps of Gram-negative bacteria and their synergism with the outer membrane barrier. Res Microbiol (2018).

42. Alexa-Stratulat, T., Pešić, M., Gašparović, A.Č., Trougakos, I.P. & Riganti, C. What sustains the multidrug resistance phenotype beyond ABC efflux transporters? Looking beyond the tip of the iceberg. Drug Resist Updat 46, 100643 (2019).

43. Piddock, L.J.V. The 2019 Garrod Lecture: MDR efflux in Gram-negative bacteria-how understanding resistance led to a new tool for drug discovery. J Antimicrob Chemother 74, 3128–3134 (2019).

44. Ricci, V., Tzakas, P., Buckley, A. & Piddock, L.J. Ciprofloxacin-resistant *Salmonella enterica* serovar Typhimurium strains are difficult to select in the absence of AcrB and TolC. Antimicrob Agents Chemother 50, 38–42 (2006).

45. Blair, J.M.A., Webber, M.A., Baylay, A.J., Ogbolu, D.O. & Piddock, L.J.V. Molecular mechanisms of antibiotic resistance. Nat Rev Microbiol 13, 42–51 (2015).

46. Darbani, B., Kell, D.B. & Borodina, I. Energetic evolution of cellular transportomes BMC Genomics 19, 418 (2018).

47. Jindal, S., Yang, L., Day, P.J. & Kell, D.B. Involvement of multiple influx and efflux transporters in the accumulation of cationic fluorescent dyes by *Escherichia coli*. BMC Microbiol 19, 195; also bioRxiv 603688v603681 (2019).

48. Grixti, J., O’Hagan, S., Day, P.J. & Kell, D.B. Enhancing drug efficacy and therapeutic index through cheminformatics-based selection of small molecule binary weapons that improve transporter-mediated targeting: a cytotoxicity system based on gemcitabine. Front Pharmacol 8, 155 (2017).

49. Klipp, E., Herwig, R., Kowald, A., Wierling, C. & Lehrach, H. Systems biology in practice: concepts, implementation and clinical application. (Wiley/VCH, Berlin; 2005).

50. Alon, U. An introduction to systems biology: design principles of biological circuits. (Chapman and Hall/CRC, London; 2006).

51. Kell, D.B. & Knowles, J.D. in System modeling in cellular biology: from concepts to nuts and bolts. (eds. Z. Szallasi, J. Stelling & V. Periwal) 3–18 (MIT Press, Cambridge; 2006).

52. Kell, D.B. Metabolomics, modelling and machine learning in systems biology: towards an understanding of the languages of cells. The 2005 Theodor Bücher lecture. FEBS J 273, 873–894 (2006).

53. Palsson, B.Ø. Systems biology: simulation of dynamics network states. (Cambridge University Press, Cambridge; 2011).

54. Mendes, P., Oliver, S.G. & Kell, D.B. Fitting transporter activities to cellular drug concentrations and fluxes: why the bumblebee can fly. Trends Pharmacol Sci 36, 710–723 (2015).

55. Bergmann, F.T. et al. COPASI and its applications in biotechnology. J Biotechnol 261, 215–220 (2017).

56. Hoops, S. et al. COPASI: a COmplex PAthway SImulator. Bioinformatics 22, 3067–3074 (2006).

57. Mendes, P. et al. Computational modeling of biochemical networks using COPASI. Methods Mol Biol 500, 17–59 (2009).

58. Hucka, M. et al. The systems biology markup language (SBML): a medium for representation and exchange of biochemical network models. Bioinformatics 19, 524–531 (2003).

59. Hucka, M. et al. The Systems Biology Markup Language (SBML): Language Specification for Level 3 Version 1 Core. J Integr Bioinform 12, 266 (2015).

60. Hucka, M. et al. The Systems Biology Markup Language (SBML): Language Specification for Level 3 Version 2 Core Release 2. J Integr Bioinform 16 (2019).

61. Iyer, R., Ferrari, A., Rijnbrand, R. & Erwin, A.L. A fluorescent microplate assay quantifies bacterial efflux and demonstrates two distinct compound binding sites in AcrB. Antimicrob Agents Chemother 59, 2388–2397 (2015).

62. Volkmer, B. & Heinemann, M. Condition-dependent cell volume and concentration of *Escherichia coli* to facilitate data conversion for systems biology modeling. PLoS One 6, e23126 (2011).

63. Stock, J.B., Rauch, B. & Roseman, S. Periplasmic space in *Salmonella typhimurium* and *Escherichia coli*. J Biol Chem 252, 7850–7861 (1977).

64. Wong, F. & Amir, A. Mechanics and Dynamics of Bacterial Cell Lysis. Biophys J 116, 2378–2389 (2019).

65. Thanassi, D.G., Suh, G.S. & Nikaido, H. Role of outer membrane barrier in efflux-mediated tetracycline resistance of *Escherichia coli*. J Bacteriol 177, 998–1007 (1995).

66. Nagano, K. & Nikaido, H. Kinetic behavior of the major multidrug efflux pump AcrB of *Escherichia coli*. Proc Natl Acad Sci 106, 5854–5858 (2009).

67. Roca, I. et al. The global threat of antimicrobial resistance: science for intervention. New Microbes New Infect 6, 22–29 (2015).

68. Laxminarayan, R., Sridhar, D., Blaser, M., Wang, M. & Woolhouse, M. Achieving global targets for antimicrobial resistance. Science 353, 874–875 (2016).

69. Gelband, H. & Laxminarayan, R. Tackling antimicrobial resistance at global and local scales. Trends Microbiol 23, 524–526 (2015).

70. Andersson, D.I. & Hughes, D. Antibiotic resistance and its cost: is it possible to reverse resistance? Nat Rev Microbiol 8, 260–271 (2010).

71. Baker, S., Thomson, N., Weill, F.X. & Holt, K.E. Genomic insights into the emergence and spread of antimicrobial-resistant bacterial pathogens. Science 360, 733–738 (2018).

72. Jindal, S., Thampy, H., Day, P.J. & Kell, D.B. Very rapid flow cytometric assessment of antimicrobial susceptibility during the apparent lag phase of bacterial (re)growth Microbiology 165, 439–454 (2019).

73. Jernaes, M.W. & Steen, H.B. Staining of *Escherichia coli* for flow cytometry: influx and efflux of ethidium bromide. Cytometry 17, 302–309 (1994).

74. Walberg, M., Gaustad, P. & Steen, H.B. Rapid preparation procedure for staining of exponentially growing *P. vulgaris* cells with ethidium bromide: a flow cytometry-based study of probe uptake under various conditions. J. Microbiol. Methods 34, 49–58 (1998).

75. Cudkowicz, N.A. & Schuldiner, S. Deletion of the major *Escherichia coli* multidrug transporter AcrB reveals transporter plasticity and redundancy in bacterial cells. Plos One 14 (2019).

76. Ishii, N. et al. Multiple high-throughput analyses monitor the response of *E. coli* to perturbations. Science 316, 593–597 (2007).

77. Kell, D.B. & Westerhoff, H.V. Metabolic control theory: its role in microbiology and biotechnology. FEMS Microbiol. Rev. 39, 305–320 (1986).

78. Fell, D.A. Understanding the control of metabolism. (Portland Press, London; 1996).

79. Zampieri, M. et al. Metabolic constraints on the evolution of antibiotic resistance. Mol Syst Biol 13, 917 (2017).

80. Lanthaler, K. et al. Genome-wide assessment of the carriers involved in the cellular uptake of drugs: a model system in yeast. BMC Biol 9, 70 (2011).

81. Girardi, E. et al. A widespread role for SLC transmembrane transporters in resistance to cytotoxic drugs. bioRxiv, 726539v726531 (2019).

82. Annunziato, G. Strategies to Overcome Antimicrobial Resistance (AMR) Making Use of Non-Essential Target Inhibitors: A Review. Int J Mol Sci 20 (2019).

83. Lamut, A., Peterlin Mašič, L., Kikelj, D. & Tomašič, T. Efflux pump inhibitors of clinically relevant multidrug resistant bacteria. Med Res Rev 39, 2460–2504 (2019).

84. Grimsey, E.M. & Piddock, L.J.V. Do phenothiazines possess antimicrobial and efflux inhibitory properties? FEMS Microbiol Rev (2019).

85. Lomovskaya, O. et al. Identification and characterization of inhibitors of multidrug resistance efflux pumps in Pseudomonas aeruginosa: novel agents for combination therapy. Antimicrob Agents Chemother 45, 105–116 (2001).

